# Distal regulation, silencers and a shared combinatorial syntax are hallmarks of animal embryogenesis

**DOI:** 10.1101/2021.09.08.459542

**Authors:** Paola Cornejo-Páramo, Kathrein Roper, Sandie M Degnan, Bernard M Degnan, Emily S Wong

## Abstract

The chromatin environment plays a central role in regulating developmental gene expression in metazoans. Yet, the basal regulatory landscape of metazoan embryogenesis is unknown. Here, we generate chromatin accessibility profiles for six embryonic, plus larval and adult stages in the sponge *Amphimedon queenslandica*. These profiles are reproducible within stages, reflect histone modifications, and identify transcription factor (TF) binding sequence motifs predictive of *cis*-regulatory elements during embryogenesis in other metazoans, but not the unicellular relative *Capsaspora*. Motif analysis of chromatin accessibility profiles across *Amphimedon* embryogenesis identifies three major developmental periods. As in bilaterian embryogenesis, early development in *Amphimedon* involves activating and repressive chromatin in regions both proximal and distal to transcription start sites. Transcriptionally repressive elements (‘silencers’) are prominent during late embryogenesis. They coincide with an increase in *cis*-regulatory regions harbouring metazoan TF binding motifs, and an increase in the expression of metazoan-specific genes. Changes in chromatin state and gene expression in *Amphimedon* suggest the conservation of distal enhancers, dynamically silenced chromatin, and TF-DNA binding specificity in animal embryogenesis.

## Introduction

Embryogenesis occurs in most animals and includes fertilization, activation of the zygotic genome, cell proliferation, cell differentiation and patterning (Kalinka and Tomancak 2012). Conserved signalling controls these processes – regulatory proteins such as Hedgehog, Notch, TGF-ß and Wnt are used across metazoan development. (Adamska et al. 2007; Larroux et al. 2008; Degnan et al. 2015; Hanks et al. 1998). Despite this conservation, embryogenesis varies markedly between and within phyla, suggesting that changes in gene expression and regulation are the basis for animal body plan diversification (Kalinka and Tomancak 2012; Wray 2003).

The chromatin environment plays a key role in regulating complex developmental programs. Within this environment, *cis*-regulatory elements, including promoters and enhancers, orchestrate the precise gene expression patterns required for multicellular development. Via their interaction with *trans*-acting proteins, *cis*-regulatory elements modulate the chromatin accessibility landscape and define cell states, identities and developmental fates. Where promoters are primarily involved in initiating transcription, enhancers play an essential role in tuning expression in a spatiotemporal context (Long et al. 2016). The interplay between *trans* factors that bind to *cis*-regulatory sequences, especially the binding of pioneer transcription factors (TFs) at enhancer regions to open up chromatin, governs gene regulatory programs by modulating gene expression (Zeitlinger 2020).

Sponges (Poriferans) are widely considered the earliest branching extant animal phyla. Their genome and gene repertoires are congruous to other animals, even though their body plan is simple – sponges do not have a nervous system, muscle cells or a gut. Their regulatory genome is complex and animal-like, with an extensive repertoire of non-coding elements, including microRNAs, long non-coding RNAs and piwiRNAs (Grimson et al. 2008; Gaiti et al. 2017; Calcino et al. 2018). *Amphimedon* larva and adult show regulatory innovations, including distal regulatory elements and bivalent promoters (possessing both activation and repressive histone marks), both of which are not found in the unicellular relative *Capsaspora* (Gaiti et al. 2017; Bulger and Groudine 2011; Bernstein et al. 2006; Sebé-Pedrós et al. 2016; Fernandez-Valverde and Degnan 2016). Furthermore, despite the lack of primary sequence conservation and the absence of shared cell types, developmental enhancers in conserved microsyntenic regions in *Amphimedon* drive cell-type-specific expression at developing vertebrates (Wong et al. 2020). This last discovery suggests a *cis*-regulatory grammar arose before the divergence of sponge and vertebrate lineages some 700 million years ago, and was maintained in conserved genomic regulatory blocks, and the coevolution of TFs through their recognition motifs.

Advances in sequencing technology have enabled the mapping of the gene regulatory landscape in many animals and cell types during embryogenesis. Transcriptomic datasets compared across the development of multiple animal phyla suggest the largest divergence in gene expression occurs during mid-development (Levin et al. 2016). Post-translational histone modifications have also been developmentally profiled in several species, although this is restricted to a few stages (Gaiti et al. 2017; Daugherty et al. 2017; Domcke et al. 2020; Bogdanović et al. 2012; Jänes et al. 2018; Schwaiger et al. 2014). With the advent of transposase-accessible chromatin using sequencing (ATAC-seq) (Buenrostro et al. 2015), genome-wide profiling of chromatin accessibility across development can be undertaken using small amounts of starting material, to provide insights into the genomic environment where the transcriptional machinery operates (Sebé-Pedrós et al. 2018b; Daugherty et al. 2017; Esmaeili et al. 2020).

To study the chromatin dynamics of embryogenesis in an early-diverged metazoan, we profiled chromatin accessibility of *Amphimedon queenslandica* across eight life stages. We interrogated differentially accessible regions across developmental stages to identify the collection of *cis*-regulatory motifs underpinning *Amphimedon* embryogenesis. We use these results to test the ability of *Amphimedon* chromatin accessible sequences to predict other species’ developmental *cis*-regulatory regions using a machine-learning framework. We integrated chromatin structure and gene expression at matched life stages to characterize developmental dynamics and to infer the regulatory genome of early metazoans.

## Results

### Dense chromatin accessibility landscapes across Amphimedon embryogenesis

To investigate the genome-wide dynamics of chromatin accessibility in *Amphimedon queenslandica*, we mapped transposase-accessible chromatin by short-read sequencing (ATAC-seq) across eight life stages. *Amphimedon* is a viviparous sponge and embryonic stages occur throughout the year in brood chambers (Degnan et al. 2015). Embryogenesis is staged by the location and pattern of pigment cells in the embryo (**Fig. 1A**). Early cleavage stages are termed white stage embryos. At the white stage, blastomeres are irregular in size and shape and mixed with maternal nurse cells. The transition to a two-layer embryo is called the brown stage and is characterised by dispersed pigments. This is followed by the cloud stage, in which the pigment cells mark the anterior-posterior axis. After this, pigment cells begin to concentrate at the posterior pole. Spot and ring stages are defined by these pigment patterns and characterised by the appearance of certain cell types.

**Figure 1.**
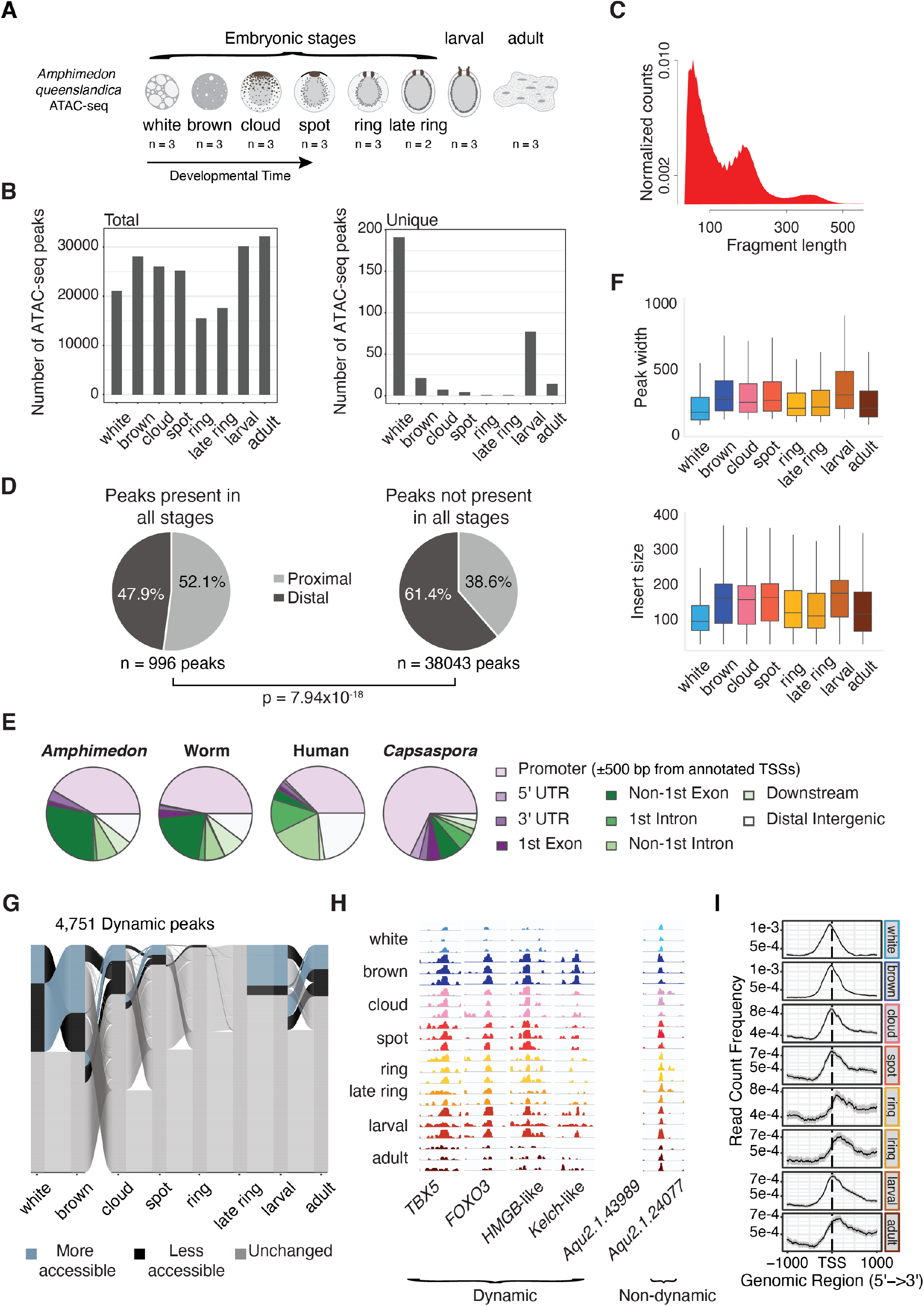
Overview of *Amphimedon cis*-regulatory regions. **(A)** *Amphimedon queenslandica* developmental stages. Number of ATAC-seq libraries for every developmental stage is shown. **(B)** Total and unique number of ATAC-seq peaks by developmental stage. Numbers for each stage calculated using the arithmetic mean across replicates. A peak must have at least a normalized count above 10 in at least one stage. Summary plot across stages required peaks with a mean count per million above 0 across all replicates for each stage. **(C)** Density plot of ATAC-seq fragment length (base pair). **(D)** Pie charts show the number of proximal (±500 bp of the TSS) and distal peaks for (i) constitutively open peaks (ii) those not accessible in all developmental stages. *χ*^2^ test was used to compute p-value. Accessible peaks across all stages: over zero normalized counts across all libraries. Peaks with varying accessibility in all stages: normalized count over 1 in 3 or more libraries. **(E)** Boxplot of ATAC-seq peak width and insert size. Numbers for each stage calculated using the arithmetic mean across replicates. **(F)** Distribution of *Amphimedon, C. elegans*, human and *Capsaspora* ATAC-seq peaks across genomic features (Daugherty et al. 2017; Buenrostro et al. 2013; Sebé-Pedrós et al. 2016). Promoter region is defined as a region within 500 bp of the TSS for all species. Downstream is defined as ≤ 300 bp of the end of a gene **(F)** Alluvial plot shows peak dynamics across life stages for peaks that change between at least one life stage (n = 4,751) (n = 808, 1422, 334, 202, 3, 0, 851 and 509 more accessible peaks for white, brown, cloud, spot, ring, late ring, larval and adult, respectively; n = 1422, 808, 693, 213, 48, 0, 202 and 563 less accessible peaks for the same stages). Differential accessibility determined by beta-binomial model (**Methods**). **(G)** Genome browser view of read coverage at selected dynamically accessible regions (*TBX5: Aqu2.1.27488; FOXO3: Aqu2.1.27411; HMGB-like: Aqu2.1.41331; Kelch-like: Aqu2.1.41157; Aqu2.1.43989*) and a selected consistently accessible peak (*Aqu2.1.24077*) across all life stages. **(H)** Chromatin accessibility read density around the TSS (within 1 kb) by life stage. Peaks were used only if at least 50% of bases overlapped across biological replicates.

We profiled individual animals from the following embryonic stages: white, brown, cloud, spot, ring, late ring embryonic stages; planktonic larval and sessile adult stages in triplicates (duplicate in late ring stage) (**Supplemental Table S1**). We identified 40,218 non-overlapping peaks (p < 1×10^-5^). These spanned 32 Mb (20%) of the *Amphimedon* genome. Median number of peaks and the average fraction of reads in peaks across libraries were 19,225 and 23%, respectively (**Supplemental Table S2**). Peak counts were strongly correlated among biological replicates (**Supplemental Fig. S1-2**). As expected, peaks were over-represented in the non-quiescent regions of the genome, based on histone marks in adults (H3K4me3, H3K27ac, H3K27me3, H3K4me1, PolII, H3K36me3) (Gaiti et al. 2017). ~64% of consensus ATAC-seq peaks matched a non-quiescent region compared to the genome background (42%) (Binomial test p = 2.1×10^-322^, OR > 2).

Adults have the greatest peak numbers, which likely reflect the complexity of cell types in adulthood, and a higher number of expressed genes (**Fig. 1B; Supplemental Table S3**). However, the greatest number of stage-specific peaks was found at the white stage. Furthermore, the typical periodicity of read fragment density, which reflects the regular positioning of nucleosomes, was not observed in the white stage in any biological replicate (**Fig. 1C; Supplemental Fig. S3**). Genes proximal to white stage-specific peaks show a 6-fold enrichment in apoptosis-related pathways (GO term ‘anoikis’, hypergeometric test, p = 1.3×10^-4^), suggesting that these peaks reflect an abundance of maternal nurse, which undergo apoptosis in the early embryo (Degnan et al. 2015; Eden et al. 2009).

While the *Amphimedon* genome is compact and gene dense (~60% of the genome is genic) (Schwaiger et al. 2014), most ATAC-seq peaks (61%) were located over ± 500 bp from the transcriptional start sites (TSS) of coding genes (**Supplemental Fig. S4**). Overall, distal peaks were more dynamically regulated while proximal peaks were more likely to be constitutively active (χ^2^ test, p = 7.9×10^-18^, OR = 1.7) (**Fig. 1D**). Accordingly, promoters tend to be constitutively accessible across cell types, while distal regulatory elements reflect cell-specific differences (Klemm et al. 2019; Bulger and Groudine 2011). Proportions of distal intergenic (10-20%) and proximal regulatory regions (40%) corresponded more closely between sponges, worms, flies and humans than the unicellular organism *Capsaspora* (Sebé-Pedrós et al. 2016) (**Fig. 1E; Supplemental Fig. S5**).

Peaks numbers were comparable to the numbers of *cis*-regulatory elements identified in other metazoans including *C. elegans* and zebrafish (Daugherty et al. 2017; Jänes et al. 2018; Bogdanović et al. 2012). Similar to fruit fly and human, peak widths ranged between 260 - 538 bp (Sebé-Pedrós et al. 2016) (**Fig. 1F; Supplemental Table S3**). Despite the rapid evolution of regulatory sequences, we found 436 proximal and 772 distal *Amphimedon* peaks aligned to the human genome with 1 bp or more overlap potentially suggesting a small degree of regulatory conservation (BLASTN E-value < 1×10^-3^). The association with the closest sponge gene revealed that these distal peaks were enriched in environmental sensing terms (hypergeometric test, FDR < 7×10^-8^; **Supplemental Table S4**).

To interrogate changes during developmental transitions, we assessed differential chromatin accessibility between stages and found that 4,751 peaks were differentially accessible between consecutive life stages (**Fig. 1G-H**). The greatest change in accessibility occurred during early embryogenesis in the transition between white and brown stages, supporting our earlier observation on white stage-specific peaks (**Fig. 1B, G**). To investigate chromatin accessibility at TSS, we mapped the density of ATAC-seq reads up and downstream of this region for each stage of development. The highest level of accessible chromatin was located immediately before the TSS in the white stage but after the TSS in the ring, late ring and adult stages (**Fig. 1I**). Furthermore, reduced accessibility downstream of the TSS was observed in the white stage, suggesting that transcription has not yet been initiated, despite accessible chromatin at the TSS, suggesting that the measured RNA in the white stage is predominantly maternally deposited. Downstream accessibility increased during embryogenesis, indicating a gradual increase in transcription in the developing embryo and the loss of maternal cells (**Fig. 1H**). Consistent with this, many genes accessible during early embryogenesis in *Drosophila*, zebrafish, mouse and human are not transcribed until later in development (Pálfy et al. 2020; Blythe and Wieschaus 2016; Lu et al. 2016; Wu et al. 2016).

### Amphimedon embryogenesis involves transcriptionally activating and repressive chromatin

Developmental cell fate decisions involve the interplay between repressive and active interactions. TF and *cis*-regulatory elements frequently repress genes (Zeitlinger 2020; Koenecke et al. 2017; Pang and Snyder 2020). Open chromatin regions can harbour *cis*-regulatory elements with activating or repressive potential (Bernstein et al. 2006; Schoenfelder et al. 2018). To identify potential activating and repressive *cis*-regulatory elements, we integrated chromatin accessibility with gene expression data. We leveraged the availability of a comprehensive set of CEL-Seq data for matched *Amphimedon* life stages (Levin et al. 2016) (n = 61 samples). Of the 10,766 expressed genes with a median count per million of 10 in at least one stage, 7451 genes (69%) have an ATAC-seq peak within 1 kb of the TSS. Most active genes were near at least one ATAC-seq peak, presumably many marking the promoter (**Fig. 2A**). Genes adjacent to several regulatory elements were associated with transcription and cell-to-cell communication (**Supplemental Fig. S6**). 31% of expressed genes were not proximal to an ATAC-seq peak at any life stage (**Fig. 2B**; shown for each stage).

**Figure 2.**
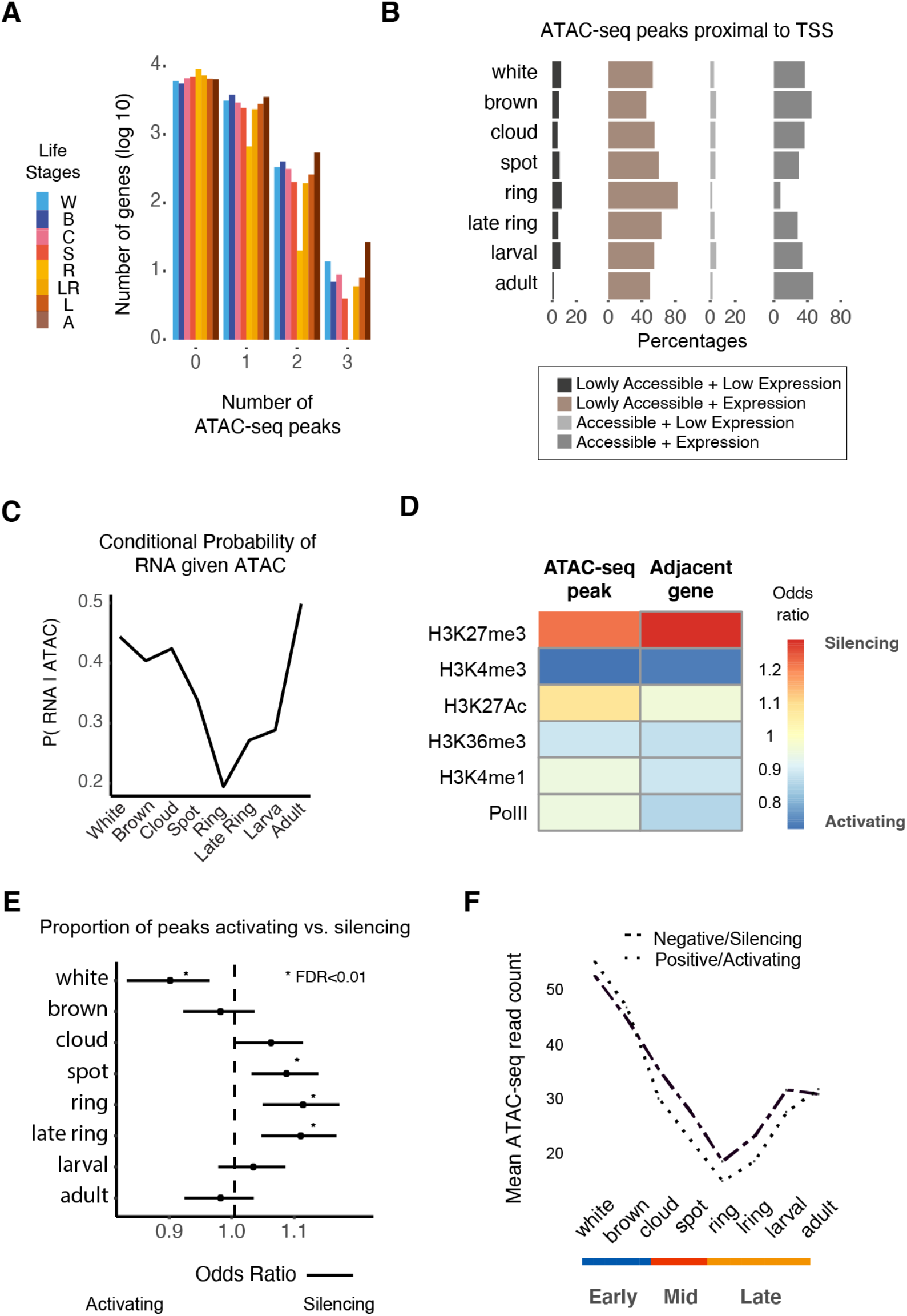
Interplay between transcription and proximal *cis*-regulatory elements. **(A)** Number of *cis*-regulatory peaks near *Amphimedon* genes (flanking 1 kb of the TSS). Number of genes was log_10_ transformed. Three genes were proximal to four *cis*-regulatory regions in late ring, larval and adult stages (1 each; not shown). **(B)** Percentage of chromatin accessible/inaccessible peaks near expressed/unexpressed genes for each life stage (flanking 1 kb of the TSS) (expressed genes were defined as those with >= 1 median cpm in stage; accessible peaks were those with >= 1 median normalized counts in every stage). **(C)** Conditional probability of gene expression given the ATAC-seq peaks accessibility. **(D)** Heatmap denotes histone mark enrichment for active and silencer peaks and their associated genes. Active and repressive peaks are defined by rank-based correlation analysis between chromatin accessibility and gene expression across time. **(E)** Forest plot shows the proportion of peaks that are active versus repressive at each life stage. Fisher’s exact tests are used to assess the significance of change relative to the total number of active and repressive peaks identified. Bars denote 95% confidence intervals. **(F)** Change in average chromatin accessibility read counts for active and silencer peaks.

To understand the interplay between accessible chromatin and transcription while accounting for peak number variation between stages resulting from variable sequencing depth, we first used conditional probability to examine the relationship between gene expression and chromatin accessibility. The probability of gene expression was calculated for each stage conditional on the probability of a proximal/promoter peak. We found that the conditional probability of gene expression, was reduced during embryogenesis at ring stages (19% compared to 40 – 50% in other stages) (**Fig. 2C**).

We integrated chromatin accessibility and gene expression data to identify activating and repressive *cis*-regulatory regions. We took a two-step approach to identify regulatory pairs first by associating peaks that were up to 1 kb from an active TSS. We used LASSO regression (Tibshirani 1996) to determine the most informative peaks and classified peaks into either repressive or activating based on the correlation of chromatin regions to the expression of genes across life stages. A *cis*-regulatory region with increased accessibility with increasing gene expression will positively correlate (termed ‘activating’). On the other hand, a *cis*-regulatory element correlated to decreased expression would have a coefficient term below zero (termed ‘repressive’, i.e. increased accessibility at these regions corresponded to lower expression across time). Where only one peak was proximal to a gene, ordinary least squares regression was used. 5,254 peaks positively correlated with gene expression and 3,686 peaks were negatively correlated.

To assess whether such assignments of activating and repressive regions were biologically meaningful, we overlapped these regions to histone marks profiled in adult *Amphimedon*. Attesting to our overall ability to distinguish between *cis*-regulatory regions with opposing regulatory function, we found a high correspondence between inferred gene activity based on histone marks and peak classification. Active chromatin marks in adults, particularly H3K4me3, were highly enriched at positively correlated peaks and genes proximal to these regions. On the other hand, the repressive polycomb-mediated H3K27me3 mark, also profiled in adults, was enriched at negatively correlated peaks, consistent with H3K27me3 regions marking silencers (Cai et al. 2021) (**Fig. 2D**). Linking this corroborating information to developmental stages, we saw a shift towards increased silencing at mid and late development, supporting our initial observations, while activating peaks were most prevalent during early embryogenesis and adulthood (**Fig. 2E-F**). In line with this, we found increased accessibility at motifs of RE1-silencing transcription factor (REST) during late development (FDR < 0.01) (**Supplemental Table S5; Methods**), where regions bound by REST have been associated with the H3K27me3 mark during the differentiation of murine neuronal cells (Arnold et al. 2013).

In summary, by integrating chromatin accessibility and gene expression our results reveal that transcriptional repression plays a crucial role in dynamically controlling *Amphimedon* development. The repressive chromatin marks, H3K9me3 and H3K27me3, are lacking in some unicellular organisms (Sebé-Pedrós et al. 2016), supporting the notion that transcriptional control of development through repressive elements is critical to the evolution of multicellularity.

### Human TF binding motifs separate developmental transitions in Amphimedon

To elucidate the dynamic changes in DNA sequence associated with developmental gene expression, we used position-weighted matrices (PWMs) to search for TF binding motifs underlying accessible chromatin. In our use of PWMs, we exploited the fact that TF gene families are deeply conserved (Kribelbauer et al. 2019; Nitta et al. 2015) and used mammalian matrices to identify known motifs. To assess motif enrichment for each time point, we combined motif alignment scores with ATAC-seq counts to measure the accessibility of each motif for each library (n = 386 PWMs) (**Supplemental Table S5; Methods**). We used a set of background peaks matched for GC content and average accessibility to assess motif enrichment.

Unsupervised clustering of motif accessibility scores clustered the libraries into three major groups recapitulating the groupings by developmental trajectory based on both peak counts and gene expression (**Fig. 3A-D**). We identified 85, 17 and 74 up-regulated differential accessible motifs at early, mid and late developmental stages, respectively (adjusted p-value < 0.05), with greatest changes, both in terms of significance and total number of motifs, occurring in early embryogenesis (**Fig. 3E-F**). Top motifs at early embryonic stages were associated with TFs linked to stem and cancer cell states, including MAX, E2F4 and Kruppel like factors. Motifs enriched during mid-embryogenesis included RREB1, tumour suppressor TP53 and glucocorticoid receptor NR3C1. Late development showed enrichment for the FOS::JUN dimer, MEIS and ISL motifs. To explore regulatory motifs underlying accessibility dynamics that did not correspond to known motifs, we performed *de novo* searches for 8-mers and identified differentially enriched accessible regions between sponge developmental stages (**Supplemental Table S6; Supplemental Fig. S7**). We identified 32,896 enriched 8-mers across consensus peaks relative to a random background set of matched GC content and average number of fragments across all stages (**Methods**). Of these, 12,523, 6,080, and 7,277 8-mers were differentially accessible between early, mid and late development, respectively (FDR < 0.05). We searched for similarities of the top six 8-mers for each stage, ranked by statistical significance (n = 18 motifs), against JASPAR PWMs. Only six of these 18 8-mers showed discernible similarities to known PWMs (q-value < 0.5; **Supplemental Table S7**).

**Figure 3.**
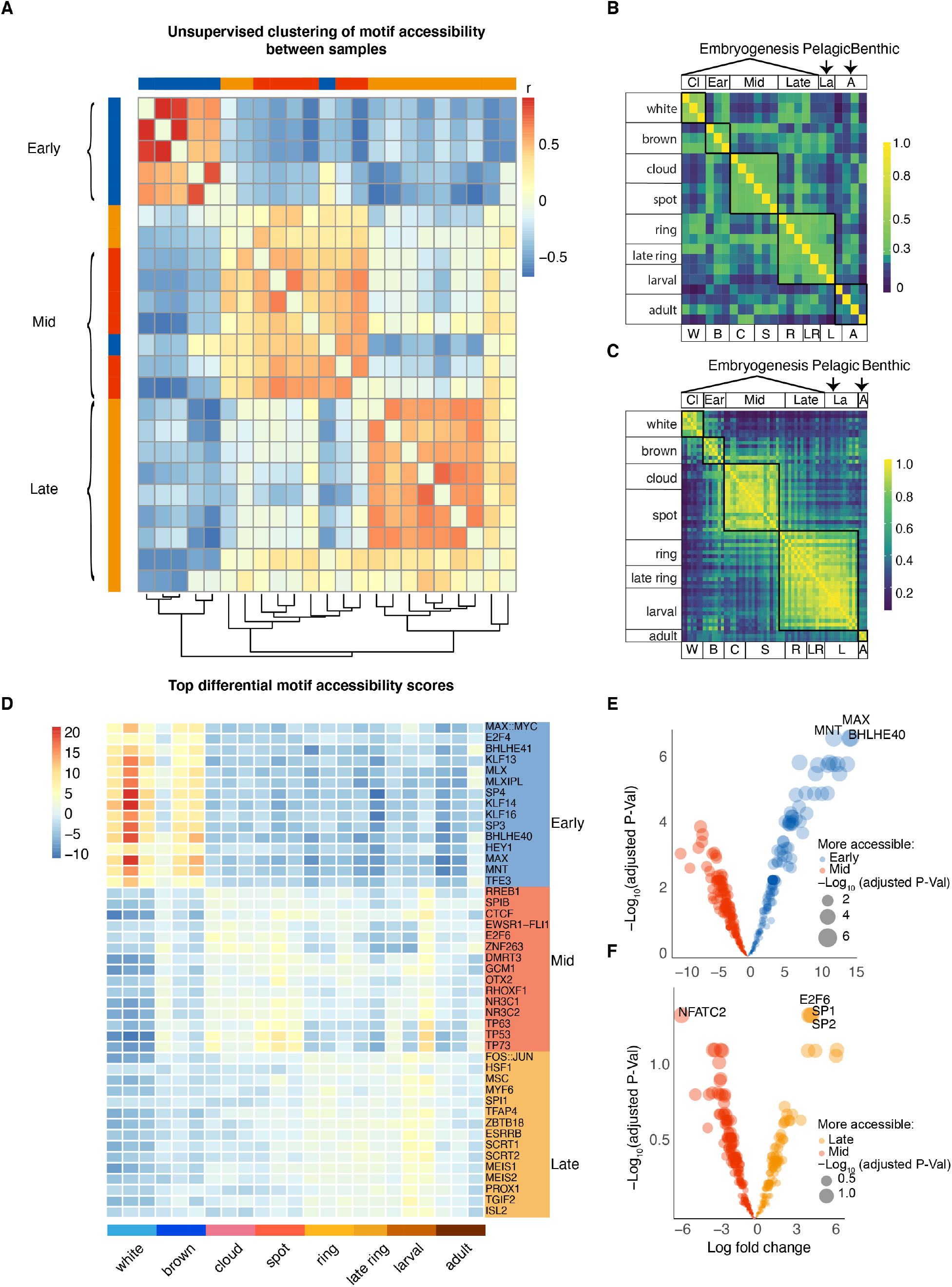
Motif analyses define three major *Amphimedon* developmental transitions. **(A)** Heatmap of Pearson’s correlation for each library based on TF motif deviation scores. Hierarchical clustering of rows and columns was performed (**Methods**). **(B)** Hierarchical clustered heatmap of Pearson’s correlation of most variable 2000 peaks based on log_10_ transformed ATAC-seq read counts across life stages. **(C)** Hierarchical clustered heatmap of most variable 1000 genes based on expression (counts per million) across life stage. **(D)** Heatmap of top differentially enriched motifs (right) based on motif accessibility where heatmap values represent motif deviation z-score (**Methods**). **(E)** Volcano plot of differentially enriched motifs between early and mid-stages **(F)** Volcano plot of differentially enriched motifs between mid and late stages. Each coloured dot represents a motif, and the size of the dot is relative to the -log_10_(q-value). x-axis is the log fold change. Early: white and brown, Mid: cloud, spot, Late: ring, late ring and larval.

TFs are known to cooperatively or competitively interact in binding to DNA, and enhancers that harbour different TF binding sites show more activity than those with a single binding site (Smith et al. 2013; Zinzen et al. 2009). We examined the motif co-localization among the top ten most differentially accessible JASPAR motifs at each *Amphimedon* developmental stage in a pairwise manner. TF motifs that co-localized at early-stage peaks include MNT & SP4 and SP4 & BHLHE40 (z-score > 10; **Supplemental Table S8**). A candidate *Amphimedon* ortholog to *MNT*, a member of the *MYC/MAX/MAD* network, (*Aqu2.1.41999*) and two *SP4*-like orthologs (*Aqu2.1.26963, Aqu2.1.26964*) have been annotated. Although, an *Amphimedon* ortholog of human *BHLHE40*, which regulates circadian rhythm and cell differentiation, has not been identified, bHLHb genes including *ARNTL*, a core component of the circadian clock in mammals that interacts with BHLHE40 (*Aqu2.1.29954, Aqu2.1.05065*) are annotated.

We further examined TF cooperatively and antagonism by calculating the correlation coefficient of motif pairs across life stages. In contrast to testing for motif co-occurrence above, we measured the correlation of motif accessibility across development. A positive correlation suggests the proteins are active at the same developmental stage and may function in similar pathways or networks. A negative correlation coefficient suggests the binding proteins do not function in the same molecular pathway. As expected, for the top ten most differentially accessible motifs at each stage, motif pairs were generally strongly positively correlated, suggesting their cognate TFs were active at the same stage and may cooperate in similar molecular processes (**Supplemental Table S9**). For example, the YY1 and KLF13/14 motifs were both highly accessible during early embryogenesis (Pearson’s rho = 0.9). Incidentally, these motifs also tend to co-occur in the same peak suggesting their cognate proteins may interact (OR = 2.30). In contrast, a strong negative correlation was apparent between YY1 and TP53 (Pearson’s rho = −0.7), where YY1 was most accessible during mid-embryogenesis when TP53 was low. Consistent with this, YY1 negatively regulates TP53 and has dual activator and repressor in other animals (Sui et al. 2004).

In summary, we find chromatin accessible regions harbour specific combinations of TF motifs in a developmental stage-dependent manner. Remarkably, these relationships at *Amphimedon* TFs can be inferred using human TFs profiles revealing a potential deep conservation of TF binding-DNA specificity.

### Upregulation of metazoan TFs during late embryogenesis

In zebrafish and fly, gene expression time-series show strong phylogenetic signatures reflecting gene age during development (Domazet-Lošo and Tautz 2010). We asked whether a similar evolutionary pattern for gene regulation is present in *Amphimedon*. To this end, we applied an evolutionary approach combining information on the phylogenetic age of *Amphimedon* genes with expression and chromatin accessibility data. Expression values were grouped by the age of the associated gene and normalized to total expression (**Methods**). Genes of eukaryotic-origin initiated early in *Amphimedon* development, while metazoan-specific genes dominated the expression profile as development progressed (**Fig. 4A-D**). In the analysis, we used 4,967 expressed sponge genes that mapped to human (Treefam); 2,853 were of eukaryotic origin, 1,003 of metazoan origin and 1,110 originated earlier at the opisthokonts (Li et al. 2006). An increase in the expression of metazoan-specific genes as development progressed has also been observed in fly and zebrafish (Domazet-Lošo and Tautz 2010).

**Figure 4.**
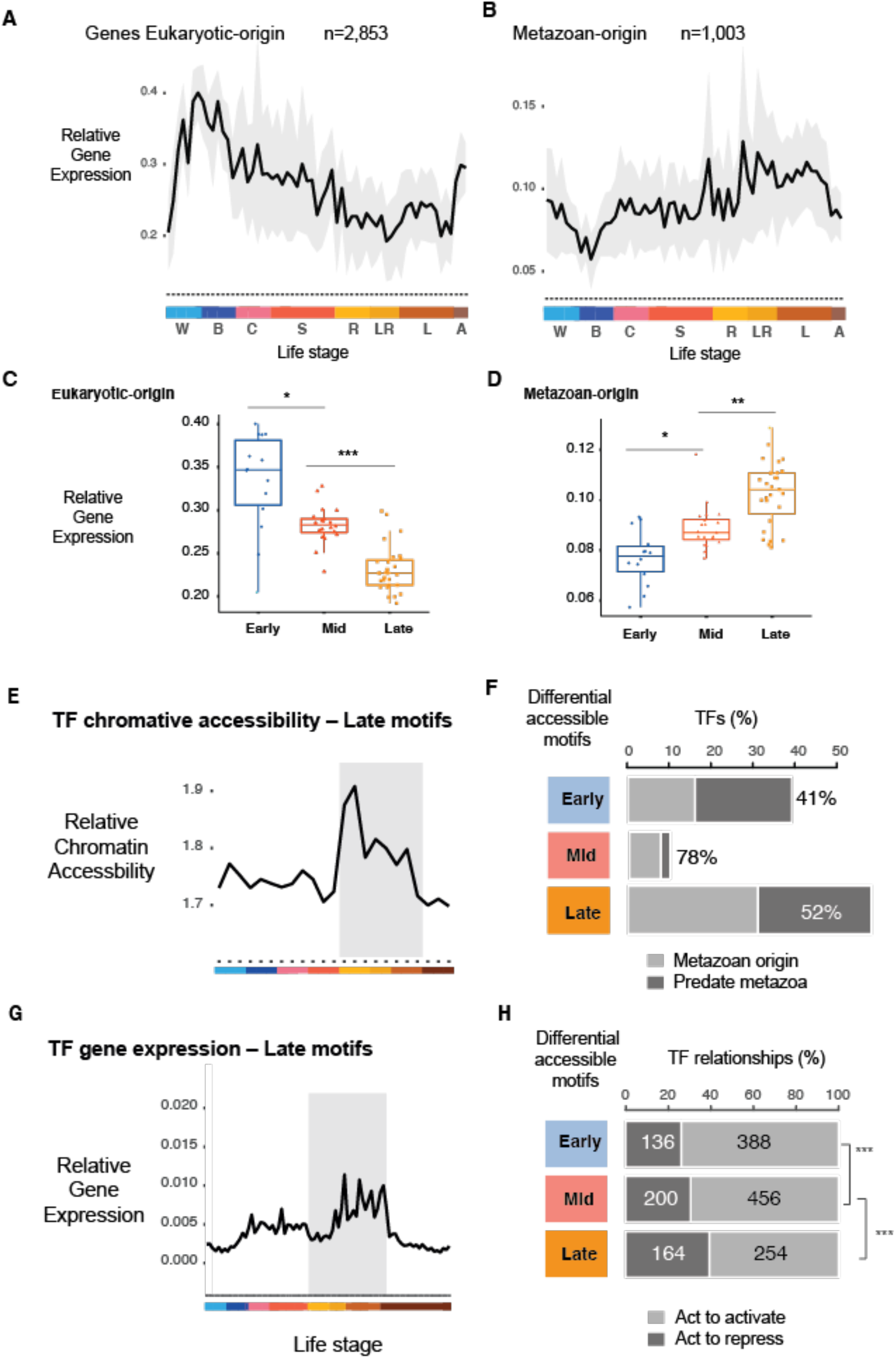
Metazoan TF motifs are enriched in late development. **(A)** Relative expression values for sponge genes that can be traced to a eukaryotic ancestor. Gray area denotes 95% CI as determined by bootstrapping. Colour denotes life stages. **(B)** Relative expression values for *Amphimedon* genes traced to the metazoan stem. **(C)** Relative chromatin accessibility values at TFs whose binding motifs are enriched in late *Amphimedon* development. ‘*’ p < 0.01, ‘**’ p < 0.001 and ‘***’ p < 0.0001 (Mann-Whitney *U* test). **(D)** Relative expression values for *Amphimedon* TFs genes enriched in late *Amphimedon* development. Significance values indicated as in **(C)**. **(E)** Bar plots show the number of TFs whose motifs are differentially enriched for each stage, grouped to whether the TF originated in metazoan stem versus those that predate metazoan (based on the TreeFam *Amphimedon* vs. human comparison). Percentage denotes TFs from the metazoan stem. **(F)** Bar plots depict the number of activating versus repressive genes based on human database TTRUST (Han et al. 2018). Genes for each stage are TFs associated to stage through differential motif analyses. Numbers denote the number of unique interactions found for that gene in the activatory or repressive category. *** indicates p-value significance from Fisher’s exact tests - early versus late: p = 2×10^-5^, and mid versus late: p = 4×10^-3^.

In a similar manner, using our ATAC-seq data, we further tested whether this trajectory was reflected by chromatin accessibility at proximal *cis*-regulatory elements. Here, high scores implied increased accessibility at the promoters of younger genes, while low values reflected the accessibility at promoters of more ancient genes. Like gene expression, the *cis*-regulatory elements of metazoan genes became increasingly accessible in late development (**Fig. 4E**). Late developmental peaks were highly enriched for motifs from metazoan genes compared to genes whose origin predate metazoans (eukaryotic or opisthokont) (Fisher’s Exact test p = 2×10^-8^, OR = 4.6) (**Fig. 4F**). In contrast, early developmental peaks were enriched for motifs of ancient/eukaryote-specific genes (**Fig. 4F**, OR = 1.9). TFs linked to motifs we previously identified at late developmental peaks were also more highly expressed during late development than at other stages (**Fig. 4G**).

Focusing on these late developmental motifs, we further sought to determine the mode of action (activating/repressive) of effector to target gene/s. We took the 15 most differentially accessible TF motifs between late development and other time points. We inferred target genes by connecting these TFs to genes via the occurrence of the motifs at open chromatin. Over 50% of the peaks containing these motifs can be associated with an expressed gene (within 1 kb of the TSS). We associated these genes using TTRUST, a curated database of knowledge on activating or repressive genes using orthology information between *Amphimedon* and human (Han et al. 2018). TFs of motifs enriched in late development were significantly more likely to show a repressive mode of action than early and mid-stage TFs (**Fig. 4H**; Fisher’s Exact test, p = 6.4×10^-5^, OR = 1.6).

In summary, an increase in metazoan gene expression as embryogenesis progresses appears characteristic of animal embryonic development. We found a similar evolutionary signature based on chromatin accessible promoter regions. Motifs at chromatin accessible region during late development corresponded to highly expressed metazoan TFs enriched for repressive function, revealing highly consistent findings across multimodal data (DNA sequence, chromatin accessibility, gene expression).

### Amphimedon cis-regulatory motif composition distinguishes developmental cis-regulatory elements

To understand the sequence basis for our chromatin accessible regions, we used a tree-based machine learning framework, to determine whether TF binding sequence motifs could distinguish *Amphimedon* peaks from genome-wide background (**Methods; Fig. 5A; Supplemental Fig. S8**). We constructed a positive peak set comprising of sampled ATAC-seq peaks and a negative set by randomly sampling regions of the same size from the genome (**Methods; Fig. 5A**). We generated ten balanced data sets for model training and testing, where for each data set, a matrix was constructed with rows as ATAC-seq peaks and columns as JASPAR Core motifs or the 18 developmental stage-enriched *de novo* 8-mers identified above. The matrix was populated using motif counts and used as input to the XGBoost gradient boosted decision trees algorithm (**Methods**). Results showed accuracies of ~0.65 in classifying between proximal and distal *Amphimedon* peaks (**Fig. 5B, Supplemental Fig. S9**; distal defined as >1kb upstream from TSS). Motif importance scores were highly correlated among the ten randomly subsampled balanced sets, suggesting robustness to peak selection (**Supplemental Fig. S10-11; Supplemental Table S10-13**). Proximal and distal regions showed representative differences reflecting sequence differences between promoter and distal regulatory regions (**Fig. 5C-D, Supplemental Fig. 12, Supplemental Fig. S13A**). Of the 20 most explanatory TF motifs, promoters were best explained by plant TF motifs, and these were often associated with environmental sensing. For example, the g-box motif (CACTG), present in plant promoters at light responsive genes (Shen et al. 2008) explain promoters but not distal elements (**Supplemental Fig. S13B**). In contrast, metazoan TF motifs (e.g. Optix, Deaf1, MXI1) were only predictive at distal regions (**Supplemental Fig. S13B**).

**Figure 5.**
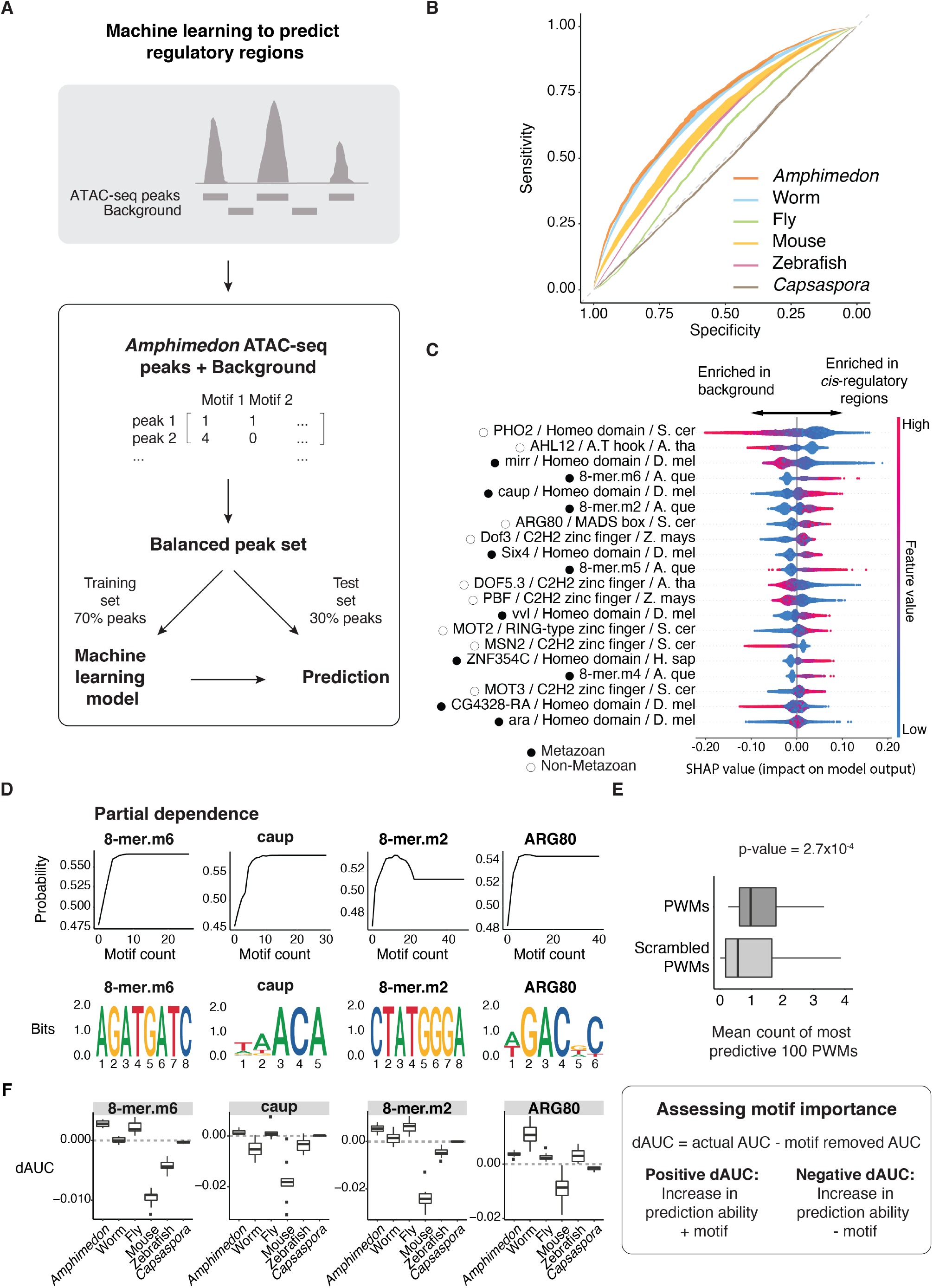
A machine-learning model trained on *Amphimedon* motifs predicts metazoan developmental *cis*-regulatory elements. **(A)** Extreme gradient boosting (XGB) machine was used on 10 balanced datasets of *Amphimedon* distal *cis*-regulatory regions and random ‘background’ peaks. 70% of peaks of every subset was used to train an XGB model. Motif counts for each peak were used predict *cis*-regulatory regions in the *Amphimedon* testing sets (30% of peaks) and in other species data **(B)** Receiver operating characteristic (ROC) curves of *cis*-regulatory regions prediction in animal species (*Amphimedon*, worm, fruit fly, mouse, zebrafish) and *Capsaspora*. The curves represent the confidence intervals (95%) calculated across the 10 models. **(C)** Relative importance score (SHAP) of most predictive known motifs and 8-mers for *Amphimedon* distal *cis*-regulatory regions. The name, class and species of motifs are indicated (S. cer = *S. cerevisiae*, A. tha = *A. thaliana*, D. mel = *D. melanogaster*, H. A. que = *A. queenslandica*, Z. mays = *Z. mays*, H. sap = *H. sapiens*). Every dot in the model represents a peak used for the training of one of the XGB models. Color represents the feature value of motifs. Metazoan and non-metazoan TFs are indicated with black filled and black outlined circles, respectively. **(D)** Top row: Partial dependence of top four most predictive PWMs of distal *cis*-regulatory regions (from **(C)**) showing the relationship between the number of instances of the motifs and the probability of a region being an actual ATAC-seq peak. Bottom row: Sequence logos of the motifs shown in **(D)** top row. **(E)** Boxplots of the mean number of instances of the top 100 most predictive actual PWMs (dark grey) and permuted PWMs (light grey), Mann-Whitney *U* test p-value shown (estimate = 0.41). Outliers removed from the plot and defined as values smaller than 1Q - 1.5×IQR or bigger than 3Q + 1.5×IQR, where “1Q” is the 1st quartile, “3Q” represents the 3rd quartile and IQR (interquartile range) is the difference between the 3Q and 1Q. **(F)** Schematic of the process to calculate dAUC as a measurement of the effect of individual motifs on distal *cis*-regulatory regions prediction ability (right); dAUC values for the top four most informative PWMs (based on **(C)**).

Motifs of key developmental metazoan TFs were significantly overrepresented at *cis*-regulatory regions, including GATA-type, homeodomain, and Forkhead domain factors (**Supplemental Table S14**). Among TF families, the TALE homeodomain superclass was overrepresented at *cis*-regulatory regions (Fisher’s Exact test, FDR = 4×10^-2^; **Supplemental Fig. S14**). This includes the MEIS1 motif which was differentially accessible during late *Amphimedon* embryogenesis (*AmqTALE, Aqu2.1.41527*) (Larroux et al. 2008).

Finally, we investigated the importance of *Amphimedon* TF binding motifs in distinguishing between metazoan and non-metazoan species. We tested whether models trained on *Amphimedon* regions could classify similar developmental regulatory elements in metazoan species including in worm, fruit fly, mouse, zebrafish and non-metazoan unicellular organism, *Capsaspora* (Sebé-Pedrós et al. 2016; Daugherty et al. 2017; Bogdanović et al. 2012; Pijuan-Sala et al. 2020; Floc’hlay et al. 2021). Negative sets were constructed controlling for *cis*-regulatory region size by sampling for size-at chromosome-matched regions from each species’ genome (**Fig. 5A, Supplemental Fig. S15A**). Models trained on motifs identified at *Amphimedon* regulatory regions could predict proximal and distal developmental *cis*-regulatory regions from the respective genome background in worm, mouse, zebrafish and fruit fly, but not *Capsaspora* (Area under ROC: 0.6-0.7 metazoan, 0.5 for *Capsaspora;* **Fig. 5B; Supplemental Fig. S9; Supplemental Table S15-16**). Comparing the number of JASPAR motifs per regulatory peak to scrambled PWMs showed that actual motifs were significantly more abundant than motifs identified using scrambled PWMs, suggesting that PWMs identified biologically important information content (**Fig. 5E; Supplemental S15B**; Mann-Whitney *U* test, p < 2.7×10^-4^; top 100 most informative PWMs). Normalizing across all datasets to adjust species-specific differences in peak widths did not change the overall findings (**Supplemental Fig. S16; Methods**). Systematically removing each motif and recalculating prediction scores for each species revealed the most informative motifs in *Amphimedon* (see **Fig. 5C**) were also informative of regulatory elements in other species (**Fig. 5F; Supplemental S17**).

In summary, we found proximal and distal *Amphimedon* regulatory regions could be distinguished based on known TF motifs. Metazoan *cis*-regulatory regions were also better identified than by chance using sequence motifs in sponge regulatory regions. Although *Amphimedon* regulatory regions were overrepresented in motifs of TFs key to metazoan development, some of the most informative overrepresented motifs have non-metazoan origin.

## Discussion

Using ATAC-seq to assay accessible chromatin in individual embryos, larvae and adults, we mapped the *cis*-regulatory dynamics during *Amphimedon* development. Chromatin accessibility is dynamically regulated across sponge embryogenesis, and changes in accessibility are concordant with changes in gene expression. Despite the compactness of the *Amphimedon* genome, the regulatory landscape appears dominated by distal (>500 bp from the TSS), rather than proximal, open chromatin regions. We identified transcriptionally repressive elements indicative of repressive enhancers (‘silencers’), which are characteristic of development in other animals (Liu et al. 2016; Pang and Snyder 2020; Koenecke et al. 2017). For instance, the repressive H3K27me3, together with the activating H3K4me3 histone modification, are highly dynamic even at the two-cell stage in mammalian embryonic development (Liu et al. 2016). Distal silencers may be unique to metazoans given the lack of H3K27me3 modifications and distal regulatory elements in the unicellular *Capsaspora* (Sebé-Pedrós et al. 2016). Their apparent functioning in sponge embryogenesis suggests gene repression is an important regulatory innovation in animal multicellularity, particularly in regulating embryogenesis.

Despite the rapid evolution of *cis*-regulatory elements, the repertoire of TF binding motif in *Amphimedon* regulatory regions is remarkably like bilaterian animals. Sponge developmental transitions were well described by human PWMs, despite many human TFs lacking one-to-one orthologs in *Amphimedon*. Our result suggests structural constrains may limit both TF family diversity and binding specificity, and thus the evolution of markedly different TF binding sites. In line with this, studies of TFs from the same structural family have been shown to recognize similar DNA sequences despite evolutionary divergence – the expanded repertoire of TFs in humans compared to fruit flies does not appear to have generally produced new TF specificities (Kribelbauer et al. 2019; Nitta et al. 2015). However, we note that our motif searches did not detect 40 bilaterian-specific motifs in *Amphimedon* (**Supplemental Table S17**). Although new TF-DNA specificities can lead to bilaterian innovations, it is also possible that other mechanisms, such as changes in the arrangement and context of shared motifs, are also important in the emergence of new gene regulatory networks involved in different body plans.

Finally, we found that a machine learning model trained with *Amphimedon* TF binding motifs could distinguish accessible chromatin regions in other metazoans, but not in the unicellular *Capsaspora*. This suggests a divergence in the composition of regulatory sequences that can be traced back to the evolutionary divergence of metazoans and non-animals. Notably, some of the most informative TF binding motifs were enriched in our genome-wide background suggesting natural selection or neutral genome evolutionary processes in driving regional sequence composition at chromatin accessible regions. In certain cases, over-represented 8-mers in regulatory chromatin regions showed a superior ability to distinguish developmental enhancers between metazoans compared to known motifs, suggesting that there may be additional but yet uncharacterized sequences for determining genome accessibility during development (**Fig. 5C; Supplemental Table S10, S12, S14**). Due to their low sequence similarity to previously characterized motifs, these sequences may influence other determinants of chromatin accessibility rather than bind canonical TFs. For example, motifs influencing DNA shape, nucleosome positioning, and DNA methylation make significant contributions to enhancer grammar (Soufi et al. 2015; Levo et al. 2015; Barozzi et al. 2014; Domcke et al. 2015).

In summary, by profiling the chromatin landscape from an early metazoan lineage, we show that the regulatory landscape of metazoan embryonic development consists of dynamic and abundant distal non-genic regions that contain shared developmental *cis*-regulatory motifs. We suggest that these deeply conserved motifs cooperatively modulate gene expression and build regulatory circuits that are reused and rewired in the evolutionary acquisition of morphological diversity. However, how divergences of *cis*-regulatory elements, their motif context, and the specificity of TF binding contribute to new gene regulatory networks remain to be fully investigated. We anticipate that advances in single-cell multimodal data and statistical/machine learning, will enable the discovery and comparative analysis of gene regulatory processes during cell and developmental evolution.

## Methods

### Temporal profiles of *Amphimedon* chromatin accessibility

Adult *Amphimedon queenslandica* were collected from Shark Bay, Heron Island, Great Barrier Reef, Queensland Australia (23°26’37.92” S, 151°55’8.81” E) under Marine Parks permit number G16/38120.1 Sponges were transferred to a closed aquaria system at the University of Queensland, Brisbane, Australia and maintained as previously described (Degnan et al. 2008). Adult brood chambers were dissected, and embryos were staged according to (Adamska et al. 2007). Larvae naturally released from sponges were collected prior to dissecting brood chambers (Leys et al. 2008). Cell suspensions from adult sponges were generated by cutting away externally facing tissue, avoiding any potential contaminating tissues and washing three times in 0.2 μm filtered Ca+ and Mg+ free artificial seawater (CMFSW) to try to reduce any potential epiphytes. However, we acknowledge that this tissue may contain allogeneic microorganisms. Cleaned adult tissue was passed through 60 μm nylon mesh to generate a single cell suspension (Sebé-Pedrós et al. 2018a).

Dissected individual embryos were placed in FSW and loaded into a P1000 pipette tip, which had been cut and fitted with sealed 60 μm mesh at the end. This allowed a small volume of CMFSW to be used to push the embryo through the mesh using a hand-held pipette and the resulting cell suspension to be collected into a 1.7mL centrifuge tube. To collect any tissue stuck on the mesh, the filter tip was then placed in 1 x Trypsin (Sigma-Aldrich)/FSW solution and placed at 37°C for 10 min. This was washed twice with FSW and added to the initial cell suspension. The filter was visually examined under a dissecting microscope to ensure no tissue remained. The above procedure was performed with larvae and the resulting cell suspensions manually counted with a hemocytometer following trypan blue exclusion staining. A single larva was estimated to have approximately 35,000 cells. Cell suspensions from adult tissues were also counted using trypan blue staining, and an equivalent amount of cells were used to make adult libraries.

Cell suspensions of approximately 35,000 cells from individual embryos and adult samples were spin at 500xg for 5 min and ATAC-seq libraries were made according to (Buenrostro et al. 2013). Libraries using the unique primers were amplified for < 7 additional cycles. The quality of each individual library was assessed using a Bioanalyzer High-Sensitivity DNA Analysis kit (Agilent). Equal volumes of libraries from three individual replicates for each stage were then pooled and run again on the Bioanalyser. Because libraries appeared to have a large concentration of free primer, we incorporated an additional purification step by adding a 1.8x concentration of AMPure beads (Beckman Coulter), washed twice with 70% ethanol and eluted in 40 μL water. A 30 nM-pooled library was prepared for sequencing.

The ATAC-seq library pool was quantified on the Agilent Bioanalyzer with the High Sensitivity DNA kit (Agilent Technologies, 4067-4626). The pool is assessed by qPCR using the KAPA Library Quantification Kit - Illumina/Universal (KAPA Biosystems, KK4824) combined with the Life Technologies Via 7 real-time PCR instrument. Pool QC and sequencing were performed at the Institute for Molecular Bioscience Sequencing Facility (University of Queensland) using the Illumina NextSeq 500 (NextSeq control software v2.0.2/ Real-Time Analysis v2.4.11). The library pool was diluted and denatured according to the standard NextSeq protocol and sequenced to generate paired-end 76 bp reads using a 150 cycle NextSeq 500/550 High Output Reagent Kit v2 (Catalog # FC-404-2002). After sequencing, FASTQ files were generated using bcl2fastq2 (v2.17) (Illumina).

### ATAC-seq read alignment and peak calling

Each ATAC-seq library was sequenced to a mean depth of ~12 million reads. Reads were aligned using Bowtie 2 (v2.3.0) (Langmead and Salzberg 2012) to the Aqu1 genome with Aqu2.1 gene annotations. Correctly mapped paired reads and those above a MAPQ value of 10 were retained using SAMtools ‘view’ (v1.6) (Li et al. 2009) for peak calling. Peak calling was performed using MACS2 (Zhang et al. 2008, 2) using the ‘callpeak’ command, a genome size of 1.2×10^-8^, and in -BAMPE mode where insert sizes of read pairs were inferred using alignment results. A set of consensus variable-sized but non-overlapping peaks was created for downstream analysis. Peaks counts reflect the number reads mapping at each peak. To remove technical compositional biases that can manifest from variable read coverage, we scaled each library based on the expected mean counts prior to analysis. Normalization for composition biases was performed using size factors calculated with the genomic mean for each peak across all libraries using the R package ‘DESeq2’ (Love et al. 2014; R Core Team 2020). To select the most robust peaks for downstream analysis, we only kept those peaks with ten or more normalized counts in at least three libraries. BEDTools (v2.28.0) was used to calculate FRiP scores (Quinlan and Hall 2010). Irreproducible Discovery Rates (IDR), were calculated following ENCODE (Li et al. 2011). We used conditional probabilities to understand the relationship between expression and chromatin accessibility to account for potential technical differences in peak number between stages. We used the arithmetic mean to average the number of counts across biological replicates for each life stage for summary figures that require associating peaks to gene expression.

### Differential accessibility analysis

We performed differential accessibility analyses of peaks using R packages ‘edgeR’ and ‘voom/limma’ (Robinson et al. 2010; Ritchie et al. 2015; R Core Team 2020). Normalization for composition biases was performed using factors calculated as described above. Peaks with low counts were removed from the analysis by filtering out peaks with less than one normalized count per million reads mapped in less than four samples—our design matrix tests for differences between adjacent life stages. The ‘voom’ function was used to calculate precision weights to remove heteroscedasticity from the data. Counts were modelled by ‘lmFit’ using empirical Bayes moderation to improve the precision of peak variability. A log fold change cut-off of 1 and FDR < 0.05 was used to define significantly changed peaks.

### Integrative correlative analysis with gene expression

To understand how accessible chromatin and transcription were linked during development, we associated genes with proximal ATAC-seq peaks located in the vicinity of TSSs (1000 bases up and downstream). In linear regression across life stages, we use ATAC-seq peaks as independent variables and gene expression data as the dependent variable to identify activating and repressive *cis*-regulatory regions. If multiple peaks are adjacent to a TSS, a LASSO regression (Tibshirani 1996) was performed to determine the most informative peak. Otherwise, ordinary least squares regression was used. The direction of association was used to class peak-gene relationship as either activating or repressive, with a positive coefficient indicative of activating and a negative coefficient suggestive of the binding of factors repressing expression.

### Gene evolutionary history analysis

To examine the connection between development and gene phylogeny, we partition expressed genes (at least 3 CEL-Seq libraries with count > 10) by their estimated evolutionary age using Treefam (v9)(Li et al. 2006), where gene orthology was assigned by phylogenetic analysis. Genes were placed into the following phylogenetic clades: Eukaryota, Opisthokont and Metazoan, reflecting the oldest clade where founders of the gene can be traced to, where possible based on orthology assignment. For each phylogenetic group and across developmental datasets, we calculate a relative measure of gene expression by taking the sum of all counts of genes classified to the same evolutionary origin, and then normalizing this by the sum of total expression values across all genes. This value was bootstrapped 500 times to generate a confidence interval.

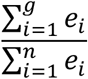

where *g* represents the genes (*i*) in the evolutionary clade and *n* represents all the genes (i) expressed in the library.

### Motif analysis

Motif analyses using both known and *de novo* sequence motif were performed using the R package ‘chromVAR’ (v3.11) (Schep et al. 2017; R Core Team 2020). The consensus set of non-overlapping peaks was used. Known human motifs from JASPAR 2016 database (Mathelier et al. 2016) were used with the default cut-off of motif calling of 5×10^-5^. For each library, an accessibility score was calculated for each motif. This is the dot product across peaks of the motif score – calculated by the match of the motif position weight matrix (PWM) and the peak accessibility score – derived from the normalized read count. Technical biases including GC content and average background accessibility, were controlled using matched sampled peaks of similar properties using the ‘getBackgroundPeaks’ function in chromVAR. A deviation score was computed, reflecting the accessibility of peaks with that motif by subtracting the expectation based on background (Schep et al. 2017). Comparison of the deviation score between three broad developmental stages (early, mid-and late) used moderated *t*-tests with false discovery rate (FDR) p-value adjustments for multiple testing (Benjamini and Hochberg 1995). Using chromVAR, we also characterized *de novo* sequence motifs, *k*-mers of length 8, within chromatin accessible regions. Deviation scores for *k*-mers are computed and compared between developmental stages using the ‘differentialDeviations’ function and a two-sided *t*-test with FDR adjustments. The top ten most deviated JASPAR motifs and top six *k*-mers for each developmental stage were used for downstream analyses. TF co-accessibility analysis was done using the ‘getAnnotationSynergy’ function in the chromVAR package, which assigns a z-score to each motif pairing reflecting variability in peak accessibility of peaks containing both motifs compared to a background sample containing only one of the two motifs. To further elucidate the relationship between motifs at accessible peaks, we examined peaks where the motif pairs did not co-locate. To do this, we calculated the Pearson’s correlation coefficient of the normalized accessibility (deviation score) across samples for each pairwise comparison, using the function ‘getAnnotationCorrelation’.

### Machine learning model

The extreme gradient boosting (XGBoost) machine learning algorithm using sequential decision trees (Chen and Guestrin 2016) was trained to distinguish between actual peaks versus randomly sampled non-overlapping regions of matched number and peak width (distal peaks were separated based on > 1kb upstream of the TSS). 10,000 *Amphimedon* peaks and 10,000 non-overlapping random peaks were sampled from the *Amphimedon* genome using the function shuffle of BEDTools (v2.27.1, - noOverlapping and -excl, the last option is used to avoid selecting background peaks that overlap with the actual *cis*-regulatory regions). Input into the algorithm is a matrix of counts of motif instances for every peak, where the positive and background sets were coded in a binary manner (actual ATAC-seq peaks = 1, random peaks = 0). Motif counts are determined using the annotatePeaks.pl function from HOMER (v4.11) (Heinz et al. 2010).

The *Amphimedon* data matrix was then split into a training dataset and a test dataset (70% and 30 % of the peaks, respectively) using R package ‘rsample’ (version 0.0.5) (Kuhn et al. 2019). Non-variable columns were removed from the training data. We train the XGM model using the *Amphimedon* training dataset using the following parameters: eta = 0.01, max_depth = 6, nround = 60000, subsample = 0.5, nfold = 10, colsample_bytree = 0.5, objective = “reg:squarederror” and early_stopping_rounds = 50 (function “xgboost” from R package ‘xgboost’ v0.90.0.2) (Chen et al. 2019), where ‘eta’ is the learning rate, ‘max_depth’ is the maximum depth of a tree, ‘nround’ represent the maximum number of boosting iterations, ‘subsample’ is the subsample ratio of training instances, ‘nfold’ is the number of random partitions of the training data, ‘colsample_bytree’ is the ratio of columns sampled when each tree is created. The XGB model is trained to minimize the ‘objective’ function and it will stop if this value does not decrease in ‘early_stopping_rounds’ rounds.

Convergence has been achieved when error did not decrease after 50 iterations. Probabilities were calculated using function ‘predict’ (type = “response”) in the R package ‘stats’ (v3.6.1). Predictions of *cis*-regulatory regions from other species (worm, fly, mouse, zebrafish and *Capsaspora*) were done in a similar way. To account for differences in peak widths across species, we trained an XGB model normalising for peak widths by dividing each count by the peak size (bp) and then multiplying by 10,000. ROC curves were generated using the function ‘roc’ from R package “pROC” (v1.16.2) (Sing et al. 2005). A threshold of 0.5 was used to transform the raw predicted probabilities into predicted classes to calculate accuracy. SHAP values were calculated using the count matrix used to train the XGB model and the model produced by xgboost (Lundberg et al. 2020).

Zebrafish (*Danio rerio*) distal regulatory elements were defined by ChIP-seq peaks of H3K4me1 excluding regions overlapping H3K4me3 regions and the TSS. The data spans four developmental stages: dome (1878 peaks), 80% epiboly (23748 peaks), 24 hpf (23419 peaks) and 48 hpf (15388 peaks) (Bogdanović et al. 2012). We sued *Drosophila melanogaster* ATAC-seq data from three developmental stages (2-4, 6-8, 10-12 hours post egg-laying) (Floc’hlay et al. 2021). Consensus peaks > 20 reads are used (7241 peaks in total). Consensus ATAC-seq peaks from *Capsaspora owczarzak* profiled from three developmental stages (filopodiated amoeba, aggregative multicellular stage and cystic stage) (11927 peaks) were used (Sebé-Pedrós et al. 2016). Mouse and worm *cis*-regulatory regions were from (Pijuan-Sala et al. 2020; Daugherty et al. 2017), respectively. Worm ATAC-seq data spans 3 developmental stages (Early Embryo, Larval stage 3 and Young Adult, n = 55432 total unique peaks). Mouse scATAC-seq peaks were filtered to select only those accessible in at least 5% of the cells profiled (corresponding to mouse embryos at 8.25 days post-fertilization) (Pijuan-Sala et al. 2020). Background peaks for worm, fly, zebrafish, mouse and *Capsaspora* selected like in sponge (option -chrom).

## Data access

All sequence data generated in this study have been submitted to ArrayExpress (https://www.ebi.ac.uk/arrayexpress/) under accession number E-MTAB-10203.

## Competing interest statement

The authors declare that they have no competing interests.

## Acknowledgements

We thank Ozren Bogdanovic for useful discussions. PC was supported by a UNSW International Postgraduate Award PhD scholarship. SMD was supported by ARC funding DP170102353. BMD was supported by ARC funding DP160100573 and FL110100044. ESW was supported by ARC funding DE160100755, DP200100250 and Advanced Queensland Maternity Fund.

## Notes

### Competing Interest Statement

The authors have declared no competing interest.

